# Ultraviolet B Induces Cutaneous Lupus Erythematosus by Triggering a Ribotoxic-Stress-Response-Dependent MIF-p38 Feedback Loop in Keratinocytes

**DOI:** 10.1101/2024.12.16.628598

**Authors:** Chipeng Guo, Siweier Luo, Jigang Luo, Haoran Lv, Yating Zhang, Le Wang, Liangchun Wang, Tao Liu, Yiming Zhou

**Affiliations:** Department of Dermatology, Sun Yat-sen Memorial Hospital, Sun Yat-sen University; Guangzhous, 510120, China; Basic and Translational Medical Research Center, Sun Yat-sen Memorial Hospital, Sun Yat-sen University; Guangzhou, 510120, China; Medical College of Acu-Moxi and Rehabilitation, Guangzhou University of Chinese Medicine; Guangzhou, 510006, China; Department of Nephrology, The First Affiliated Hospital, Sun Yat-sen University; Guangzhou, 510080, China

**Author notes:** EMAIL ADDRESSES: Chipeng Guo, Siweier Luo, Jigang Luo, Haoran Lv, Yating Zhang, Le Wang, Liangchun Wang. Yi Xiao https://orcid.org/0009-0003-4286-5408.

## Abstract

Ultraviolet B (UVB) is one of the most common triggers of cutaneous lupus erythematosus (CLE), but the mechanism is still unclear. Here, we discovered that the expression of macrophage migration inhibitory factor (MIF), a multipotent cytokine, was significantly increased in keratinocytes of lupus patients in scRNA-seq data and clinical samples. We further discovered that UVB exposure induced MIF releasing from lupus keratinocytes and contributed to the tissue remodeling and inflammation by regulating the expression of matrix metalloproteinases (MMPs) and pro-inflammatory cytokines in keratinocytes and fibroblasts. Mechanistically, UVB exposure triggered a ribotoxic stress response (RSR) in lupus keratinocytes, which was featured by the increased level of phospho-ZAKα. The RSR induced MIF releasing through the activation of p38 and GSDMD cleavage. The released MIF could further promote the activation of p38 through its receptor CD74, forming a positive feedback loop triggered by UVB exposure in lupus keratinocytes. Intradermal injection of a Mif-shRNA AAV significantly improved the skin lesions in lupus mice. To further investigate the potentials of MIF as a therapeutic target, we developed a microneedle patch capable of intradermal administration of a MIF inhibitor, which effectively protected lupus mice from skin lesions induced by UVB exposure. In conclusion, our findings suggest a novel mechanism by which UVB exposure contributes to the development of CLE by triggering a positive feedback loop of MIF-p38 in lupus keratinocytes. Our results also demonstrate the translational potentials of microneedle patch targeting MIF in the treatment of CLE.

## INTRODUCTION

Systemic lupus erythematosus (SLE) is an autoimmune disease with multiple organs involvement, which seriously threatens human health ^1,2^ and causes enormous economic burdens for society ^3^. Skin lesions of SLE, also named as cutaneous lupus erythematosus, were one the most frequent manifestations and often occurred as the earliest symptom ^2,4,5^. Phenotypes of tissue remodeling and inflammation can be found in CLE lesions ^2^, accompanied with increased expression of MMPs (e.g., MMP2 and MMP9) ^6^ and pro-inflammatory cytokines (e.g., TNFA, IL1B, and IL6) ^7^. Considering the refractory nature, frequent recurrence, disfiguring changes, and potential risk of systemic damages in CLE, developing new targets and therapies of CLE are crucial.

UVB is one of the most important triggering factors for CLE, of which this phenomenon is known as photosensitivity ^2^. To date, the damage of UVB for the disease has been widely recognized, reflected by the facts that photosensitivity is one of the diagnostic criteria for SLE ^8^, and sun protection is highly recommended for the SLE patients ^9^. Nevertheless, the pathological mechanisms of photosensitivity are still unknown ^4,10,11^. A study suggested that UVB has a wavelength of 280-320 nm, and its penetration depth is restricted in the epidermal layer ^11^. As the dominant cell type in the epidermis, keratinocytes could be the target cells of UVB and drive the photosensitivity of CLE ^12^.

Macrophage migration inhibitory factor (MIF) is a pleiotropic cytokine that is closely related to the development of SLE ^13,14^. Not only are the gene polymorphisms of MIF associated with SLE susceptibility ^15–18^, but also the plasma concentration of MIF are positively correlated with the severity of lupus nephritis ^19^. In skin tissues, immune cell- and keratinocyte-derived MIF has been demonstrated to play an important role in tissue remodeling and inflammation ^20,21^. However, the mechanism by which UVB regulates the release of MIF remains unclear.

In this study, using scRNA-seq analysis and clinical samples, we discovered that an increased population of MIF+ keratinocyte clusters in CLE patients. UVB exposure significantly promoted MIF releasing from lupus keratinocytes and subsequently resulted in the tissue remodeling and inflammation by increasing the expression of matrix metalloproteinases (MMPs) and pro-inflammatory cytokines in autocrine and paracrine manners. Mechanistically, UVB exposure induced MIF releasing by triggering the ribotoxic stress response (RSR) in lupus keratinocytes, characterized by the activation of ZAKα/p38 signaling pathway and GSDMD cleavage. In addition, we demonstrated that keratinocyte-derived MIF promoted the phosphorylation of p38 through the receptor CD74, thereby forming a MIF-p38 positive feedback loop in lupus keratinocytes. Intradermal injection of a MIF-shRNA AAV protected lupus mice from skin lesions induced by UVB exposure. To further investigate the therapeutic potentials of MIF in CLE, we developed a microneedle patch capable of intradermal administration of MIF inhibitor ISO-1, which significantly alleviated the UVB-induced skin lesions in lupus mice. In short, we discovered a novel mechanism that UVB exposure triggers a positive feedback loop of MIF-p38 in lupus keratinocytes and leads to the development of CLE. Successful translation of microneedle patches targeting MIF may offer a novel treatment for CLE patients in the future.

## RESULTS

### scRNA-seq analysis identified keratinocyte clusters with increased levels of MIF in lupus patients

The scRNA-seq data were downloaded from GEO with the accession number GSE186476. scRNA-seq data from 14 normal control (NC), 7 pairs of lupus lesional (Lupus Les) and non-lesional (Lupus NonLes) skins were included for the analysis. Dimension reduction using Uniform Manifold Approximation and Projection (UMAP) were performed and the cells were divided into 26 clusters according to their transcriptomic profiles in an unsupervised manner (**Fig. S1A**). Each cluster was annotated with their marker genes and the primary cell types of skin tissues were captured and marked, including keratinocytes, fibroblasts, melanocytes, endothelial cells, smooth muscle cells, mast cells, T cells, and myeloid cells (**Fig. S1B**). Analysis results indicated that MIF was widely expressed by a wide variety of cell types, among which keratinocytes were one of the major producers of MIF (**Fig. S1C**). Interestingly, other pro-inflammatory cytokines such as IFNG, IFNK, TNFA, IL1B, or IL6 were hardly detected in keratinocytes, indicating that MIF was a dominant cytokine in keratinocytes (**Fig. S1D**). To explore the expression levels of MIF among these three groups, we next divided keratinocytes into 9 subclusters based on the top 3 marker genes of each cluster (**Fig. S1E**). The feature plots showed that there were two unique subclusters (subcluster 3 and 7) in lupus patients (**Fig. S1F**). In addition, the expression levels of MIF in these two lupus-specific subclusters were higher in both lupus non-lesional and lesional skins (**Fig. S1G**). Collectively, the scRNA-seq analysis results suggested that keratinocytes may be one of the major cell types producing MIF in lupus skins.

### Keratinocyte-derived MIF resulted in skin tissue remodeling and inflammation

To validate the scRNA-seq analysis results, we performed immunohistochemistry (IHC) staining of skin tissues from normal controls (NC), dermatomyositis (DM), lichen sclerosus et atrophicus (LSA), and CLE patients. The IHC staining results indicated that MIF was strongly expressed in the epidermis of CLE compared with that of NC group (**Fig. 1A**). Meanwhile, the expression of the MIF receptor CD74 was notably elevated in the lupus epidermis (**Fig. 1B**). To further elucidate the expression pattern of MIF in skin cells, we co-stained MIF and its receptor CD74 with the keratinocyte marker keratin 14 (KRT14) and fibroblast marker vimentin (VIM) in the lesional skins of CLE. The co-staining results showed that MIF and CD74 were highly expressed in keratinocytes, while some of CD74 was also expressed in fibroblasts (**Fig. 1C**). The qPCR results indicated that the expression levels of MIF signaling molecules (MIF, CD74, and CD44), tissue remodeling molecules (COL I, MMP2, and MMP9), and inflammation molecules (IL1B, TNFA, IFNG, and MX1) were significantly upregulated in lesional skins of CLE (**Fig. 1D**).

**Fig. 1.**
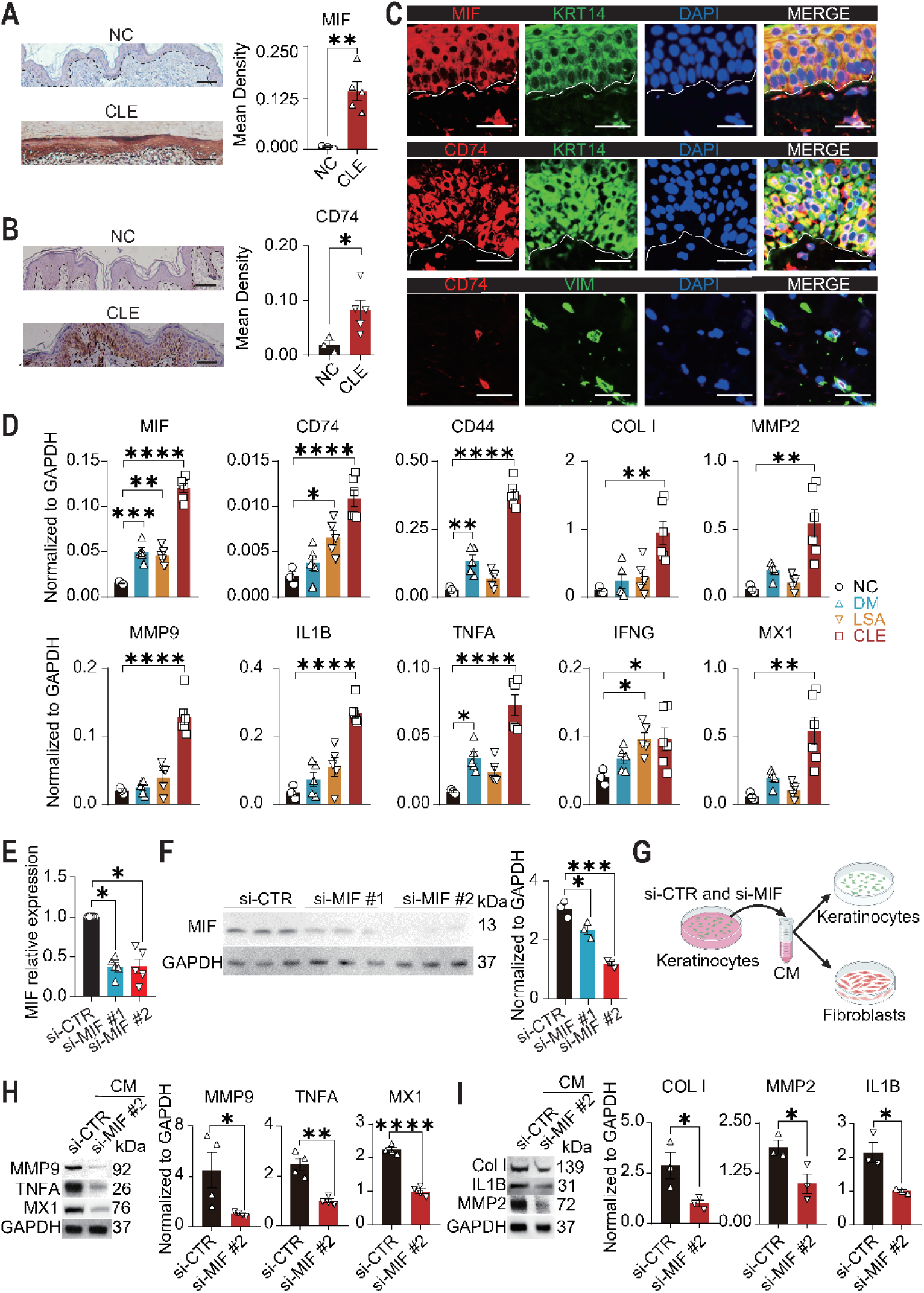
Keratinocyte-derived MIF contributed to the tissue remodeling and inflammation in CLE. **(A)** Representative images and quantitative results of immunohistochemical staining of MIF in skin tissues from normal control (NC, N = 3) and cutaneous lupus erythematosus (CLE, N = 5). **(B)** Representative images and quantitative results of immunohistochemical staining of the MIF receptor CD74 in skin tissues from NC (N = 3) and CLE (N = 5). **(C)** Representative images of immunofluorescence staining showing the co-localization of MIF and its receptor CD74 with the keratinocyte marker keratin 14 (KRT14) and fibroblast marker vimentin (VIM) in skins from CLE. MIF, CD74 and KRT14 (N = 4), CD74 and vimentin (N = 3). **(D)** The mRNA expression levels of MIF, CD74, CD44, COL1A1, MMP2, MMP9, IL1B, TNFA, IFNG, and MX1 in skin tissues from NC (N = 3), DM (N = 5), LSA (N = 5), and CLE (N = 6). **(E)** The mRNA expression levels of MIF in HaCaT cells after 24 hours transfection of a control-siRNA (si-CTR), and two MIF-siRNAs (si-MIF #1 and si-MIF #2) (N = 5 each). **(F)** The protein levels and the corresponding quantitative result of MIF in HaCaT cells after 48 hours transfection of a control-siRNA (si-CTR), and two MIF-siRNAs (N = 3 each). **(G)** Schematic diagram of the experiment design for investigating the effect of keratinocyte-derived MIF on HaCaT cells and fibroblasts. HaCaT cells and fibroblasts were cultured with the conditional medium collected from control-siRNA and MIF-siRNA transfected HaCaT cells. **(H)** Investigation of the effect of keratinocyte-derived MIF on the expression of tissue remodeling and inflammation molecules in keratinocytes. The protein levels and the corresponding quantitative results of MMP9, TNFA, and MX1 in HaCaT cells cultured with the conditional medium for 48 hours from control-siRNA and MIF-siRNA #2 transfected HaCaT cells (N = 4 each). **(I)** Investigation of the effect of keratinocyte-derived MIF on the expression of tissue remodeling and inflammation molecules in fibroblasts. The protein levels and the corresponding quantitative results of COL I, MMP2, and IL1B in fibroblasts cultured with the conditional medium for 48 hours from control-siRNA and MIF-siRNA #2 transfected HaCaT cells (N = 3 each). NC, normal controls; DM, dermatomyositis; LSA, lichen sclerosus et atrophicus; CLE, cutaneous lupus erythematosus. Data are mean ± standard error of the mean. Scale bars= 100μm. \**P* < 0.05, \*\**P* < 0.01, \*\*\**P* < 0.001, \*\*\*\**P* < 0.0001.

To examine the impact of keratinocyte-derived MIF on skin lesions, we transfected keratinocytes with MIF-siRNAs. This led to a significant decrease in the expression of MIF mRNA and protein in keratinocytes (**Fig. 1E and F**). Conditional media from keratinocytes transfected with control-siRNA and MIF-siRNA were collected and added to another keratinocytes and fibroblasts **(Fig. 1G)**. As shown, the protein levels of MMP9, TNFA, and MX1 in keratinocytes **(Fig. 1H)** and COL1A1, MMP2, and IL1B in fibroblasts **(Fig. 1I)** were significantly downregulated after the treatment with the conditional media from MIF knockdown keratinocytes. These results indicated that keratinocyte-derived MIF played an important role in skin tissue remodeling and inflammation through autocrine and paracrine manners.

### UVB exposure triggered the release of MIF from keratinocytes and exacerbated skin tissue remodeling and inflammation

Previous studies have showed that UVB exposure may regulate MIF expression ^22,23^, however, we found that there was no difference in the expression levels of MIF before and after UVB exposure at different intensities and time points (**Fig. S2**). Unexpectedly, we discovered that UVB exposure dose-dependently triggered the release of MIF from keratinocytes (**Fig. 2A**). To investigate whether UVB exposure led to tissue remodeling and inflammation via the keratinocyte-derived MIF, conditional media from untreated and UVB-exposed keratinocytes was collected and then added to another keratinocytes and fibroblasts together with DMSO and MIF inhibitor ISO-1 (**Fig. 2B**). As a result, the conditional medium from UVB-exposed keratinocytes dose-dependently increased the mRNA levels of MMP9 and TNFA in keratinocytes (**Fig. 2C**), and COL1A1 and MMP2 in fibroblasts (**Fig. 2D**). These effects were also confirmed at the protein levels in keratinocytes **(Fig. 2E-F)** and in fibroblasts (**Fig. 2G-H**). Notably, these effects could be significantly reversed by supplying the MIF inhibitor ISO-1 to the conditional medium from UVB-exposed keratinocytes **(Fig. 2E-H)** Collectively, the results indicated that UVB exposure induced the release of MIF from keratinocytes, which in turn increased the expression levels of multiple genes related to skin tissue remodeling and inflammation, therefore contributing to the development of skin lesions in CLE.

**Fig. 2.**
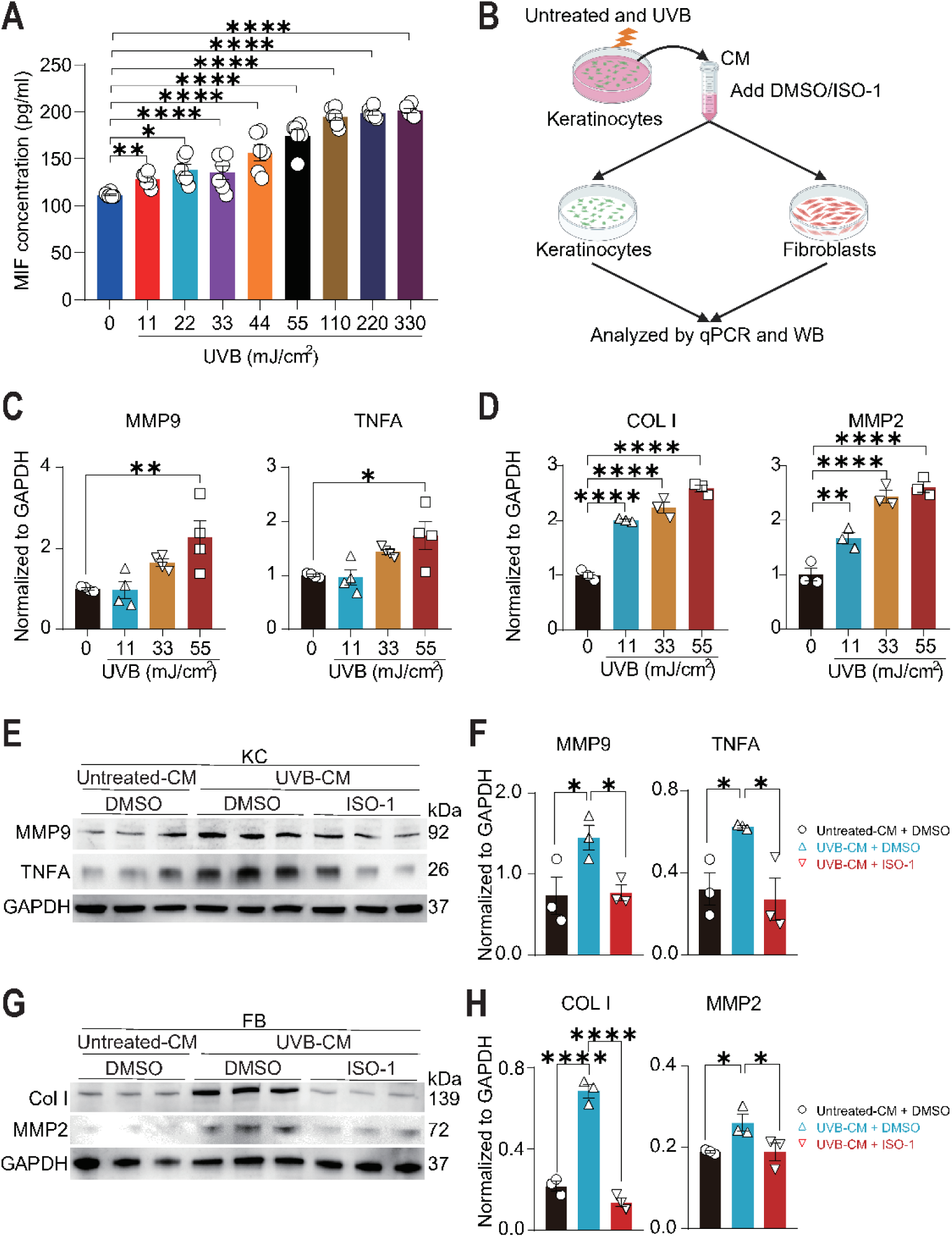
UVB exposure induced the release of MIF from keratinocytes and aggravated the tissue remodeling and inflammation *in vitro*. **(A)** ELISA determination of the levels of secreted MIF in supernatants 12 hours after UVB exposure at different intensities of 11, 22, 33, 44, 55, 110, 165, and 330 mJ/cm^2^. **(B)** Schematic diagram of the experiment design for investigating the effect of UVB-induced MIF releasing on keratinocytes and fibroblasts. HaCaT cells and fibroblasts were cultured with the conditional medium with DMSO or MIF inhibitor ISO-1, which were collected from the untreated and UVB-exposed HaCaT cells. **(C)** The mRNA levels of MMP9 and TNFA in keratinocytes cultured with the conditional medium from HaCaT cells after UVB exposure at different intensities of 11, 33, and 55 mJ/cm^2^ (N = 4 each). **(D)** The mRNA levels of COL I and MMP2 in fibroblasts cultured with the conditional medium from HaCaT cells after UVB exposure at intensities of 11, 33, and 55 mJ/cm^2^ (N = 3 each). **(E and F)** The protein levels (**E**) and quantification results (**F**) of MMP9 and TNFA in keratinocytes cultured with the conditional medium, which were collected from untreated (Untreated-CM) or UVB-exposed HaCaT cells (UVB-CM), with DMSO or MIF inhibitor ISO-1. (N = 3 each). (**G and H**) The protein levels (**G**) and quantification results (**F**) of COL1 and MMP2 in fibroblasts cultured with the conditional medium collected from the same groups, with or without ISO-1 (N = 3 each). Untreated-CM: conditional medium collected from untreated HaCaT cells; UVB-CM: conditional medium collected from UVB-exposed HaCaT cells. Data are mean ± standard error of the mean. \**P* < 0.05, \*\*\**P* < 0.001, \*\*\*\**P* < 0.0001.

### UVB exposure led to MIF releasing by activating the ribotoxic stress response in lupus keratinocytes

Recently, UVB exposure was reported to cause the ribotoxic stress response in keratinocytes, leading to the activation of ZAKα/p38 signaling pathway ^24–26^ and NLRP1/GSDMD-mediated pyroptosis ^24^. The formation of GSDMD pores could lead to cellular lysis and the release of inflammatory cytokine ^27^. Consistently, we discovered that the expression levels of RSR-related proteins, including p-ZAKα, p-p38, and GSDMD-NT in keratinocytes were significantly upregulated after the UVB exposure **(Fig. 3A-B)**. In addition, the ELISA results showed that both UVB exposure and ZAKα-activating toxin anizomycin (ANS) could significantly promote the released level of keratinocyte-derived MIF **(Fig. 3C)**. We then explored the involvements of p38 and GSDMD using the p38 inhibitor SB203580 and pyroptosis inhibitor disulfiram (DSF) in the UVB-exposed keratinocytes **(Fig. 3D-G)**. As expected, SB203580 treatment could reversed the upregulated levels of p-p38 and GSDMD-NT **(Fig. 3D-E),** while DSF could reversed the upregulation of GSDMD-NT **(Fig. 3F-G)** in keratinocytes after the UVB exposure. Of note, both SB203580 and DSF reduced the level of UVB-induced MIF releasing from keratinocytes **(Fig. 3H)**, suggesting that the p38-GSDMD associated pyroptosis pathway played an important role in the UVB-induced MIF releasing. Taken together, these results suggested that UVB exposure caused the release of MIF from keratinocytes through the activation of ZAKα-dependent RSR and its downstream p38-GSDMD-related pyroptosis pathways.

**Fig. 3.**
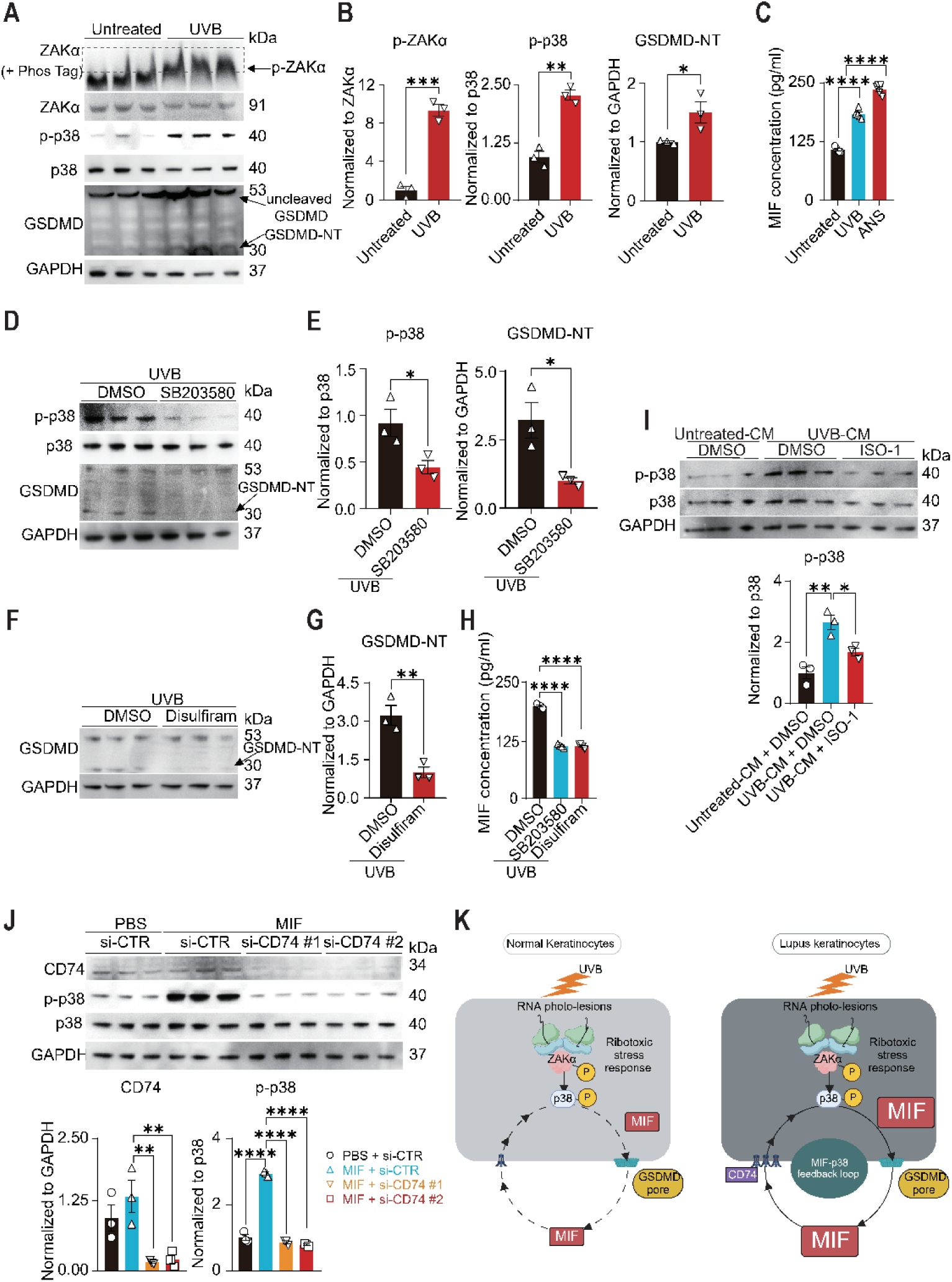
UVB exposure triggered a MIF-p38 feedback loop via the ribotoxic stress response in lupus keratinocytes. (**A and B**) The expression levels (**A**) and quantification results (**B**) of p-ZAKα, ZAKα, p-p38, p-38, and GSDMD (both uncleaved full-length and the N-terminal fragment) in HaCaT cells after UVB exposure at an intensity of 55 mJ/cm^2^ (N = 3 each). (**C**) The levels of secreted MIF in the supernatants from untreated, UVB-exposed, and ZAKα activator (ANS)-treated HaCaT cells detected by ELISA assays (N = 5 each). (**D and E**) The expression levels (**D**) and quantification results (**E**) of p-p38, p38, GSDMD, and GSDMD-NT in UVB-exposed HaCaT cells pre-treated with DMSO and p38 inhibitor SB203580 (N = 3 each). (**F and G**) The expression levels (**F**) and quantification result (**G**) of GSDMD and GSDMD-NT in UVB-exposed HaCaT cells treated with DMSO and pyroptosis inhibitor disulfiram (N = 3 each). (**H**) ELISA assay showing the level of secreted MIF in supernatants of UVB-exposed HaCaT cells treated with DMSO, SB203580, and disulfiram (N = 3 each). (**I**) The effect of ISO-1 on the expression levels of p-p38 and p38 in lupus keratinocytes treated with the conditional medium from unexposed (Untreated-CM) and UVB-exposed HaCaT cells (UVB-CM) (N = 3 each). The quantification result was showed below. (**J**) The effect of MIF on the expression levels of CD74, p-p38, and p38 in HaCaT cells transfected with a control-siRNA (si-CTR) and two CD74-siRNAs (si-CD74 #1 and #2) (N = 3 each). The quantification results were showed below. (**K**) Schematic diagram of the mechanism by which UVB triggers the p38-MIF feedback loop in lupus keratinocytes. Untreated-CM: conditional medium collected from untreated HaCaT cells; UVB-CM: conditional medium collected from UVB-exposed HaCaT cells. Data are mean ± standard error of the mean. \**P* < 0.05, \*\**P* < 0.01, \*\*\**P* < 0.001, \*\*\*\**P* < 0.0001.

### UVB exposure formed a MIF-p38 positive feedback loop in lupus keratinocytes

P38 was a key signaling hub in keratinocytes responding to external stimulation ^28,29^. To investigate the impact of secreted MIF on lupus keratinocytes, we generated lupus-like keratinocytes by transfection of endogenous nucleic acids (eNAs) to HaCaT cells as previous reported ^30,31^. We discovered that the expression levels of MIF and CD74 were significantly increased in the eNAs-transfected HaCaT cells, indicating successful generation of lupus-like keratinocytes (**Fig. S3**). We then discovered that the conditional medium from UVB-exposed keratinocytes significantly promoted the phosphorylation of p38 in lupus-like keratinocytes, of which the effect could be reduced by supplying MIF inhibitor ISO-1 (**Fig. 3I**). To confirm whether this effect was dependent on the MIF receptor CD74 in lupus keratinocytes, we knocked down CD74 in lupus-like keratinocytes using siRNA. Interestingly, the levels of p-p38 in these cells were significantly diminished upon knocking down CD74 (**Fig. 3J**), indicating that MIF may activate p38 through its receptor CD74 in an autocrine manner. These results suggested a novel mechanism of UVB effect on skin lesion of CLE, that is UVB exposure could trigger a MIF-p38 positive feedback loop via the ribotoxic stress response in lupus keratinocytes, which contributes to the skin lesions of CLE (**Fig. 3K**).

### Intradermal injection of Mif-shRNA AAV alleviated UVB-induced skin lesions of MRL/lpr mice

To investigate the effect of UVB exposure in keratinocytes *in vivo*, we applied a UVB-induced skin lesions model in lupus mice according to a previously reported method ^32^. After UVB exposure, MRL/lpr mice developed obvious skin lesions that manifested as redness and crusting **(Fig. 4A-B)**. Epidermal hyperplasia and even ulceration could be observed in UVB-exposed skins **(Fig. 4A-B)**. The IF staining results demonstrated that UVB exposure led to an upregulation of MIF expression in mouse keratinocytes **(Fig. 4C),** as well as enhanced expression levels of COL1A1, MMP9, MMP2, and TNFA in skin lesions of MRL/lpr mice **(Fig. 4D-E)**. It’s known that mouse NLRP1 is insensitive to regulation by ZAKα and p38 given the lack of the critical NLRP1 linker region in rodent NLRP1 orthologs ^24^, and UVB exposure doesn’t result in pyroptosis in mouse keratinocytes in both C57BL/6 wide-type (WT) and ZAKα-/- mice ^33^. To our surprise, UVB exposure led to stronger histopathological changes in MRL/lpr mice than both wide-type (WT) and ZAKα-/- C57BL/6 mice, meanwhile, consistent with the *in vitro* results, UVB exposure also significantly increased the levels of MIF, p-p38, and GSDMD-NT in the skin lesions **(Fig. 4F-G)**.

**Fig. 4.**
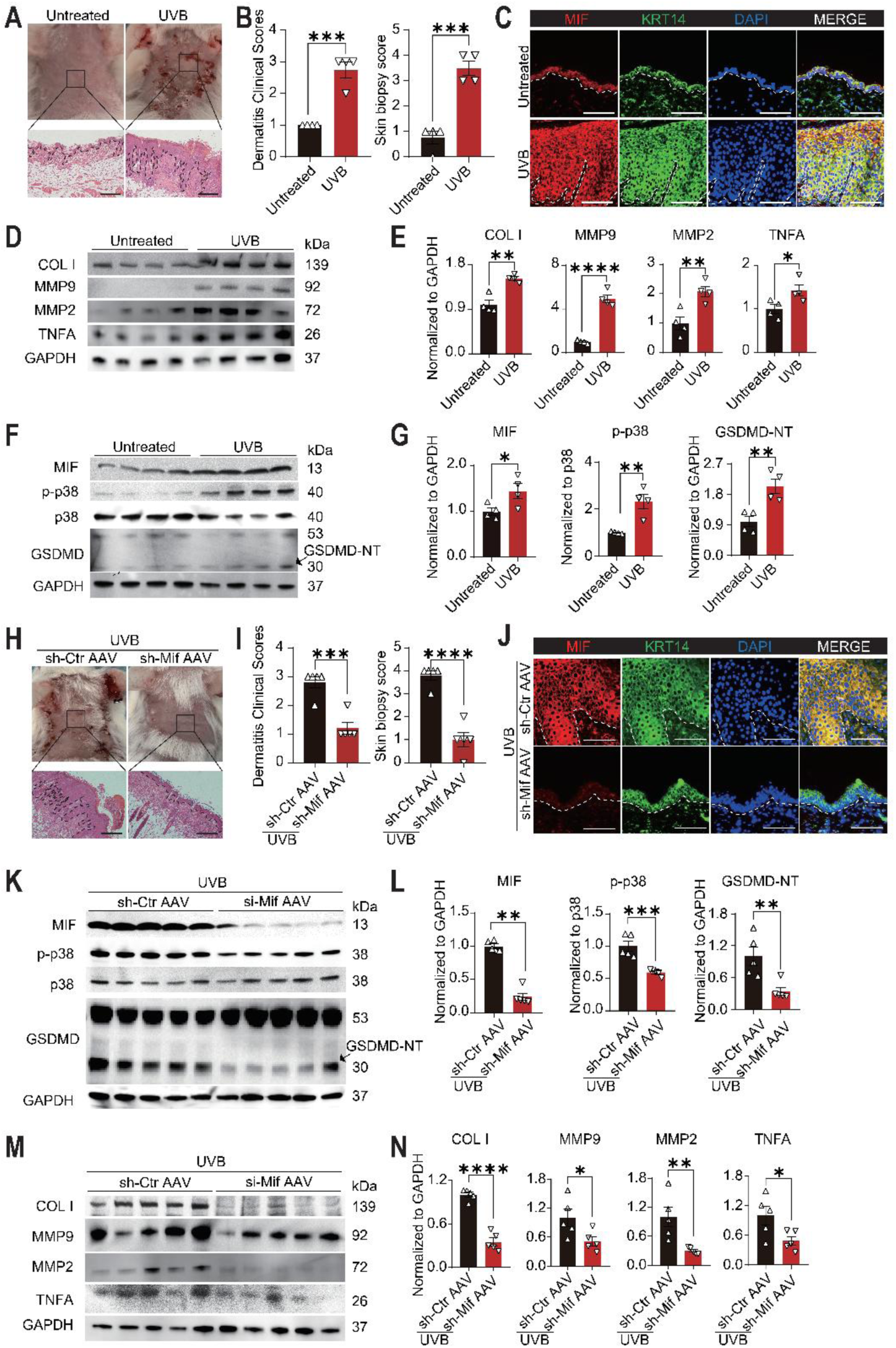
Intradermal injection of Mif-shRNA AAV alleviated UVB-induced skin lesions in MRL/lpr mice. (**A**) Representative macroscopic appearances and H&E staining images of untreated and UVB-exposed skins in MRL/lpr mice (N = 4 each). (**B**) Dermatitis clinical scores and skin biopsy scores of **(A)**. (**C**) Representative images of immunofluorescence staining showing the co-localization of MIF and KRT14 in untreated and UVB-exposed skins of MRL/lpr mice (N = 4 each). **(D and E)** The protein levels (**D**) and quantification results (**E**) of COL I, MMP9, MMP2, and TNFA in untreated and UVB-exposed skins of MRL/lpr mice (N = 4 each). **(F and G)** The protein levels (**F**) and quantification results (**G**) of MIF, p-p38, p38, GSDMD, and GSDMD-NT in untreated and UVB-exposed skins of MRL/lpr mice (N = 4 each). **(H)** Representative macroscopic appearances and H&E staining images of UVB-exposed skins in MRL/lpr mice after injection of control-shRNA AAV (sh-Ctr AAV) and Mif-shRNA AAV (sh-Mif AAV) (N = 5 each). **(I)** Dermatitis clinical scores and skin biopsy scores of **(H)**. **(J)** Representative images of immunofluorescence staining showing the co-localization of MIF and KRT14 in UVB-exposed skins of MRL/lpr mice after injection of control-shRNA AAV and Mif-shRNA AAV (N = 4 each). **(K and L)** The protein levels (**K**) and quantification results (**L**) of MIF, p-p38, p38, GSDMD, and GSDMD-NT in UVB-exposed skins of MRL/lpr mice after injection of control-shRNA AAV and Mif-shRNA AAV (N = 5 each). **(M and N)** The protein levels (**M**) and quantification results (**N**) of COL I, MMP2, MMP9, and TNFA in UVB-exposed skins of MRL/lpr mice after injection of control-shRNA AAV and Mif-shRNA AAV (N = 5 each). sh-Ctr AAV: control-shRNA AAV; sh-Mif AAV: Mif-shRNA AAV. Data are mean ± standard error of the mean. Scale bars = 100μm. \**P* < 0.05, \*\**P* < 0.01, \*\*\**P* < 0.001, \*\*\*\**P* < 0.0001.

To explore the potentials of MIF as a therapeutic target in CLE, we generated the scramble-shRNA (sh-Ctr) and Mif-shRNA (sh-Mif) AAVs, and verified the effectiveness of Mif-shRNA AAVs in reducing the expression of MIF in primary mouse keratinocytes **(Fig. S4)**. Intradermal injection of the Mif-shRNA AAVs prior to UVB exposure significantly improved the skin lesions, dermatitis clinical scores, and skin biopsy score in lupus mice **(Fig. 4H-I)**. Intradermal injection of Mif-shRNA AAVs but not scramble-shRNA AAVs significantly reduced the expression of Mif in UVB-exposed keratinocytes, suggesting that knocking down of MIF could improve the skin lesions and clinical scores in lupus mice after UVB exposure **(Fig. 4J)**. Moreover, intradermal injection of Mif-shRNA AAVs before UVB exposure significantly reduced the expression levels of activated p38 (p-p38) and cleaved GSDMD (GSDMD-NT) in keratinocytes of lupus mice **(Fig. 4K-L)**. As expected, knocking down of Mif with these AAVs reversed the UVB-induced upregulation of COL1A1, MMP9, MMP2, and TNFA in lupus mice **(Fig. 4M-N)**. Take together, we herein demonstrated that MIF could be a therapeutic target for the treatment of UVB-induced skin lesions in lupus mice.

### Blocking MIF with ISO-1-loaded microneedle patches improved UVB-induced skin lesions in MRL/lpr mice

To develop an ideal treatment targeting keratinocyte-derived MIF, we developed a microneedle patch system that could intradermally administrate a MIF inhibitor ISO-1. This microneedle patch system could increase the skin permeability of the drug and avoid excessive systemic administration **(Fig. 5A)**. The microneedles were arranged in a 10*10 array on a 10*10 mm patch and all of them were in a unified shape and distribution as shown by optical microscopy and SEM **(Fig. 5B)**. We confirmed the mechanical property of the microneedles to ensure they have enough strength for the skin insertion **(Fig. 5C)**. The compression tests revealed that the microneedles could tolerate the load force up to 8 N and to a depth of 450 μm without any fracture, suggesting that the microneedles could maintain integrity during skin insertion **(Fig. 5C)**. In addition, the methylene blue-loaded microneedle patches were inserted in the porcine skin for 5 min and an array of blue pinholes appeared at the insertion sites after removal, which indicated that the needles could penetrate deeply enough for drug delivery **(Fig. 5D)**. Of note, there was no erythema or edema of the skin at the different time points after the microneedle insertion **(Fig. S5 and Fig. 5E)**, while the degradation tests showed that the drug could be gradually released within 30 minutes after insertion **(Fig. 5F)**. These results confirmed that the microneedle patch could be an ideal material for targeting the keratinocyte-derived MIF.

**Fig. 5.**
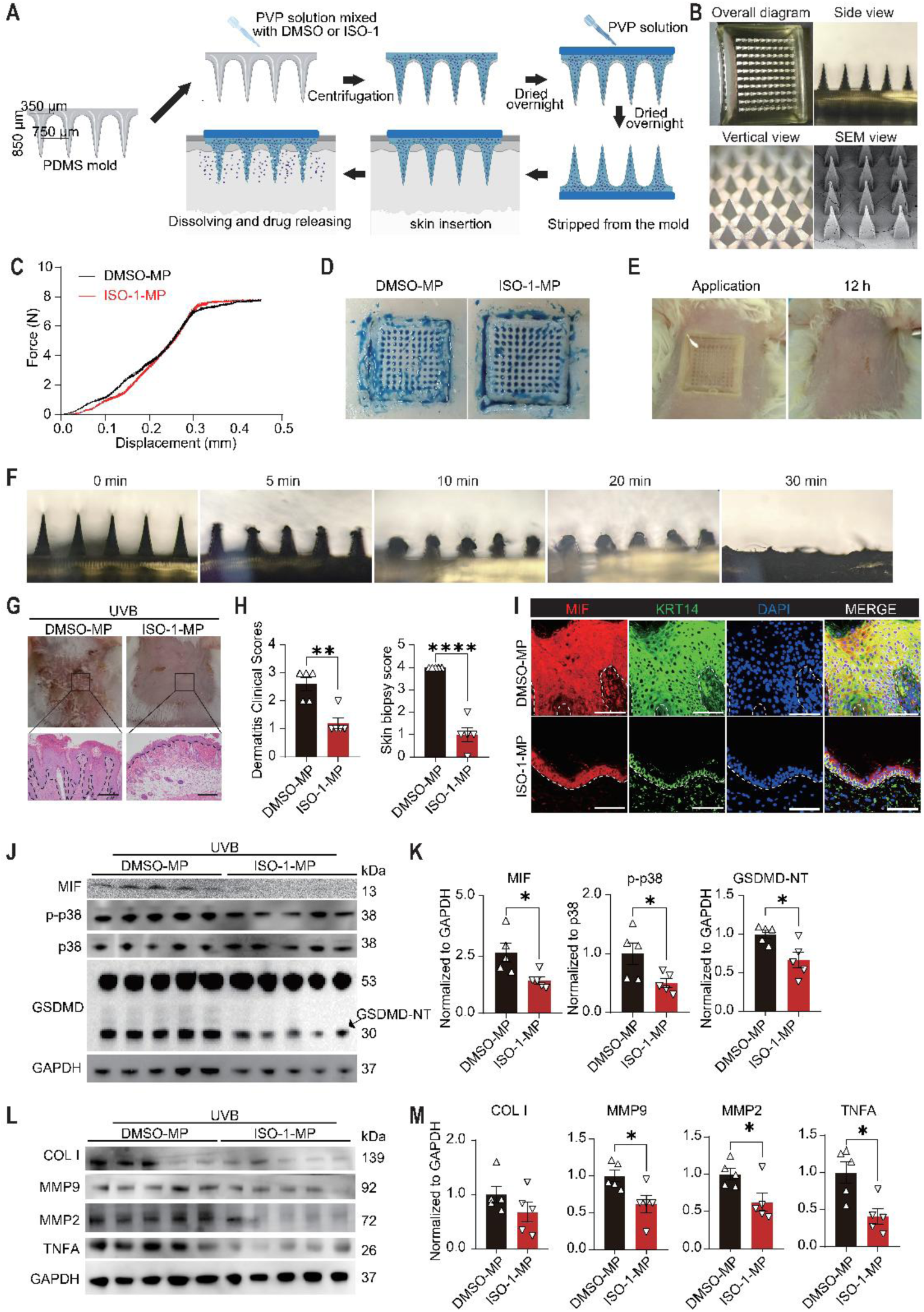
Transdermal administration of MIF inhibitor ISO-1 using microneedle patches alleviated UVB-induced skin lesions in MRL/lpr mice. **(A)** Schematic diagram showing the fabrication process of microneedle patches. **(B)** Representative stereo-microscopic images and the scanning electron microscope (SEM) image of microneedle patches. **(C)** Characterization of the mechanical strength of DMSO- and ISO-1-loaded microneedle patches. **(D)** Representative images showing the results for skin insertion tests of DMSO- and ISO-1-loaded microneedle patches stained by methylene blue. **(E)** Representative macroscopic appearances of skins in MRL/lpr mice during microneedle patch administration and 12 hours after peeling off microneedle patch. **(F)** Representative images showing the results for degradation tests of microneedle patches after being inserted in rat skins for 5, 10, 20, and 30 minutes. **(G)** Representative macroscopic appearances and H&E staining images of UVB-exposed skins in MRL/lpr mice treated with DMSO- or ISO-1-loaded microneedle patches (N = 5 each). **(H)** Dermatitis clinical scores and skin biopsy scores of **(G)**. **(I)** Representative images of immunofluorescence staining showing the co-localization of MIF and KRT14 in UVB-exposed skins of MRL/lpr mice treated with DMSO- or ISO-1-loaded microneedle patches (N = 4 each). **(J and K)** The protein levels (**J**) and quantification results (**K**) of MIF, p-p38, p38, GSDMD, and GSDMD-NT in UVB-exposed skins of MRL/lpr mice treated with DMSO- or ISO-1-loaded microneedle patches (N = 5 each). **(L and M)** The protein levels (**L**) and quantification results (**M**) of COL I, MMP2, MMP9, and TNFA in UVB-exposed skins of MRL/lpr mice treated with DMSO- or ISO-1-loaded microneedle patches (N = 5 each). DMSO-MP: microneedle patches loaded with DMSO; ISO-1-MP: microneedle patches loaded with ISO-1. Data are mean ± standard error of the mean. Scale bars = 100μm. \**P* < 0.05.

After application of ISO-1-loaded microneedle patch (ISO-1-MP), the severity of UVB-exposed skins was significantly improved, showing lower clinical scores and biopsy scores than that of DMSO-MP group **(Fig. 5G-H)**. Although the fluorescence intensity of MIF in each keratinocyte was comparable between two groups, the overall population of MIF-positive keratinocytes was significantly reduced by ISO-1-loaded microneedle patch, suggesting that this system could effectively prevent the activation of MIF-p38 feedback loop in lupus keratinocytes triggered by the UVB exposure **(Fig. 5I)**. Indeed, ISO-1-MP treatment significantly decreased the levels of MIF, p-p38, and GSDMD-NT in lupus keratinocytes **(Fig. 5J-K)**, and reversed the upregulation effect of UVB exposure on MMP9, MMP2, and TNFA **(Fig. 5L-M)**. Collectively, these results demonstrated that not only could MIF be a therapeutic target for the UVB-induced skin lesions, but also the microneedle patch system could be a promising treatment for CLE (**Fig. 6**).

**Fig. 6.**
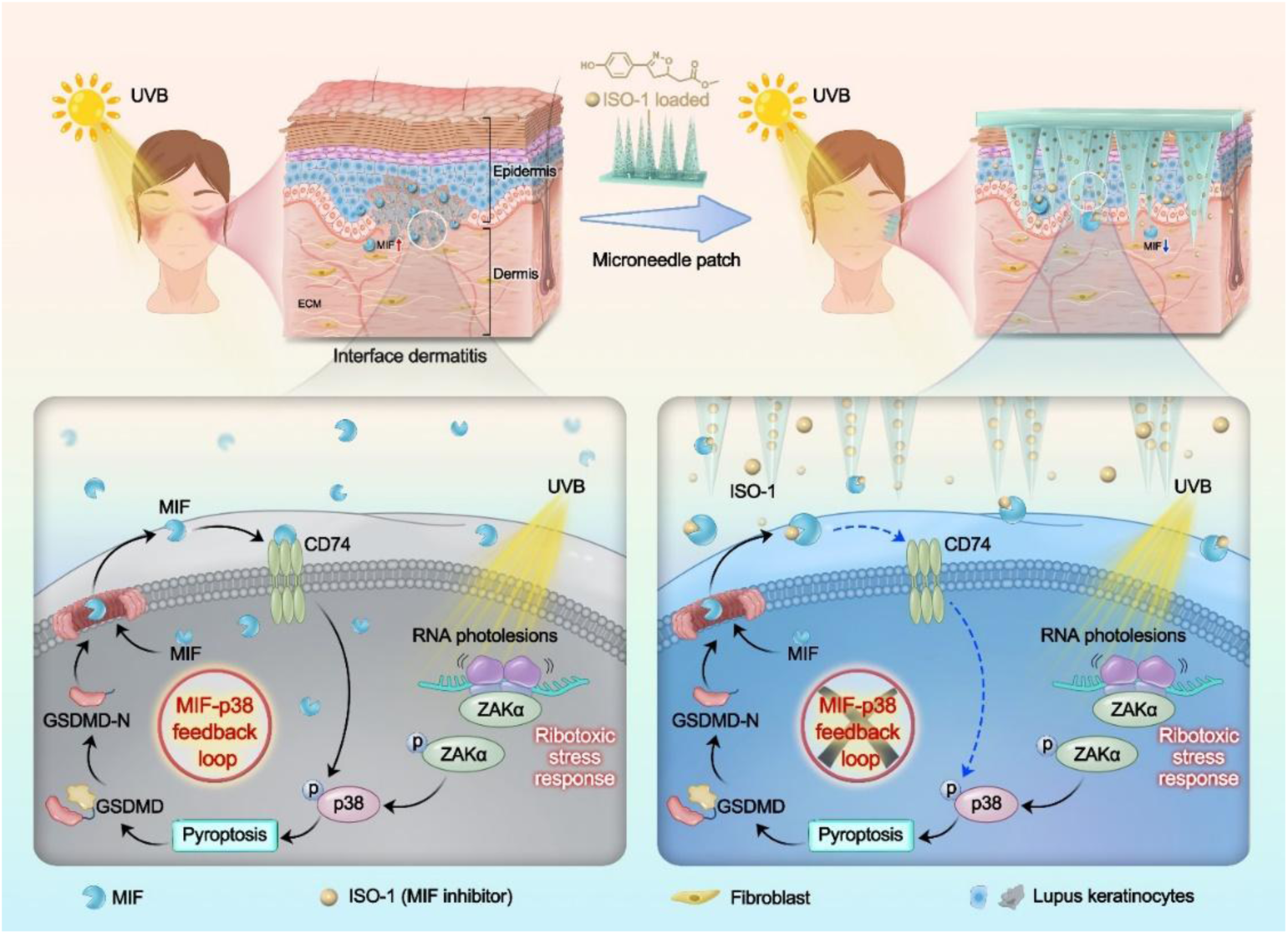
Schematic diagram of the mechanism of which UVB induces skin lesions in CLE patients by triggering a ribotoxic-stress-response that could activate a MIF-p38 positive feedback loop in lupus keratinocytes, and the effect can be reversed by transdermal hadministration of MIF inhibitor ISO-1 using microneedle patches.

## DISCUSSION

Skin lesion is one of the most common manifestations of SLE and often occurred at earlier time point ^2,4,5^. For preventing systemic involvement, early diagnosis and treatment of CLE is crucial. UVB is widely recognized as an important trigger for the disease, which could hardly be avoided in daily lives ^4,10,11^. In the context of genetic predisposition, UVB induces the occurrence of skin lesions even at relatively lower exposure intensities, and the phenomenon of photosensitivity is reflected by the lower minimal erythema dose (MED) in CLE patients ^34–36^. Previous studies suggest that UVB exposure leads to the damage and apoptosis of keratinocytes, with the insufficient clearance of the damaged and apoptotic cells leading to the accumulation of autoantigens in lupus skins, which in turn activates the innate immune and adaptive responses ^7^. Considering the challenges of treatment resistance and the potential risks for systemic involvement, it is important to clarify the molecular mechanism of the photosensitivity of CLE.

As is known, interface dermatitis is the typical histopathological changes of CLE, which is characterized by the destruction of basement membrane and the dense infiltration of immune cells in the dermis ^2^. The increased expression levels of MMPs, such as MMP2 and MMP9 ^6^, as well as inflammatory cytokines, such as type I IFNs, TNF-α, and IL-6 ^7^, are involved in the tissue remodeling and inflammation of CLE. Traditionally, the immune cell-centered model of CLE emphasizes the contributions of various immune cells including plasmacytoid dendritic cells (pDCs) ^37,38^, autoreactive T ^39,40^ and B cells ^41^ to the development of skin lesions. To date, however, the clinical results indicated that treatments targeting these immune cells are not always satisfying ^42–44^. Recent studies have demonstrated that non-immune cells, especially keratinocytes could be the initiators of autoimmunity in CLE ^42,45^. With a wavelength range of 280-320 nm, the penetration of UVB was mainly restricted at the epidermis, which suggested that keratinocyte may play an important role in photosensitivity ^11^. Studies found that lupus keratinocytes exhibited abnormal gene expression profiles ^45,46^, and released more cytokines such as TNF-α and IL-6 after the UVB exposure ^47–49^. Although different cytokines were reported to participate in the development of skin lesions, therapies targeting these molecules were still unsatisfying ^42–44^. In fact, using a scRNA-seq dataset, we found that levels of innate immunity molecules like TNF-α, IL-6 and IL-1β were barely detectable in keratinocytes. Instead, MIF was one of the highly-expressed cytokines in keratinocytes. In addition, we discovered that there were two unique keratinocyte clusters with the upregulated expression of MIF, which were found exclusively in CLE patients. MIF was a pro-inflammatory cytokine and gene polymorphism of MIF was closely related to SLE ^15–17,19^. Studies have also shown that MIF could upregulate the levels of MMPs and cytokines in keratinocytes and fibroblasts ^50–52^. However, the role of MIF in CLE is unclear.

In this study, we demonstrated the critical role of keratinocyte-derived MIF in the development of skin lesions and verified that knockdown or inhibition of MIF alleviated skin tissue remodeling and inflammation both *in vitro* and *in vivo*. We provided crucial evidence in elucidating the role of keratinocyte-derived MIF in the initiation of tissue remodeling and inflammation in CLE and the molecular mechanism of MIF mediating photosensitivity. UVB exposure was recently demonstrated to cause the ribotoxic stress response in keratinocytes, resulting in the activation of ZAKα/p38/GSDMD pathway and leading to pyroptosis ^24^. We herein demonstrated that both UVB exposure and ZAKα activator ANS led to the release of MIF from keratinocytes, which could be blocked by p38 inhibitor SB203580 and pyroptosis inhibitor disulfiram. In eNAs-transfected keratinocytes, we demonstrated that MIF could further activate p38 through the receptor CD74, which forms a MIF-p38 positive feedback loop that triggered by UVB exposure in these lupus-like keratinocytes. This positive feedback loop could also partially explain why occasionally UVB exposure leads to the severity of CLE patients. Prevention of this positive feedback loop by targeting MIF, CD74 and p38 could effectively reverse the tissue remodeling and inflammation triggered by UVB exposure.

To investigate the therapeutic potentials of MIF in treating CLE, we developed a transdermal microneedle patch system that could intradermal delivering MIF inhibitor ISO-1 to lupus mice. These microneedle patches dramatically decreased the amount of ISO-1 for targeting MIF in keratinocytes, as the systemic administration was up to 40 mg/kg daily as reported previously ^16^. As a result, ISO-1-loaded microneedle patches significantly reversed the skin lesion and clinical scores in UVB-exposed lupus mice.

Nevertheless, there are also some deficiencies in our research. Firstly, the effectiveness and safety of ISO-1-loaded microneedle patches should be further evaluated in clinics. Secondly, integration analysis of more scRNA-seq datasets is crucial to clarify the role of MIF in a larger CLE population. Thirdly, although gene polymorphism is related to the expression of MIF in SLE, we didn’t explore the contributions of environmental factors in regulating MIF gene. These limitations notwithstanding, our results not only suggest a novel mechanism of how UVB exposure triggers the pathogenesis of CLE, but also providing a potential treatment for CLE by targeting MIF using microneedle patch. Successful translation of this work may provide a novel treatment for CLE patients in the future.

## MATERIALS AND METHODS

### Experimental Design

We aimed to explore the photosensitivity of CLE and search for novel treatments for the disease in the study. The plans of the research were to (i) investigate the expression of MIF in lupus keratinocytes and detect the role of keratinocyte-derived MIF in UVB-induced skin lesions of CLE and (ii) explore the efficacy of MIF-targeted therapy in CLE through AAVs and microneedle patches. The study received approval from the Medical and Ethics Committees of Sun Yat-Sen Memorial Hospital, Sun Yat-Sen University. Written informed consent was obtained from all the subjects. All animal experiments were approved by the Institutional Animal Care and Use Committees of Sun Yat-Sen University.

### Clinical samples

The diagnosis of CLE was determined by clinical and histological evidence ^53^, while the classification criteria for SLE was based on the 2019 European League Against Rheumatism/American College of Rheumatology criteria ^9^. This study excluded patients who had both malignancy and active infection. Normal skin tissues were collected from the adjacent skin of removed-pigment nevi and all of them were UVB-unexposed. The lesional and normal tissues were age- and sex-matched. **Table S1** displayed information about the donors and the sites of tissue origin.

### Animal experiments

The 12-week-old female MRL/lpr mice were purchased from Jiangsu Huachuang Xinnuo Medical Technology Co., LTD. The mice were bred in a pathogen free (SPF) environment and maintained under a 12:12 h light–dark cycle with optimal humidity (60–80%) and temperature (22 ± 1 °C). No more than five mice were co-housed per cage.

### UVB exposure

The UVB irradiation instrument (SS-01, SIGMA, Shanghai) used in animal and cell experiments has a wavelength range of 290-320nm, with a peak value of 311 nm and an irradiation power of 11.00 mw/cm^2^. In the animal experiment, the exposure area of each mouse was located at a hair removal skin on the back, with a diameter of 1.5 cm. The exposure area was irradiated for 4 cycles, each cycle was irradiated for 5 days and rest for the subsequent 2 days. The irradiation energy was 330 mJ/cm^2^ during the former 2 cycles and 110 mJ/cm^2^ during the last 2 cycles ^32^ **(Table S2)**. After UVB exposure, the severity of skin lesions was evaluated using dermatitis clinical scores and skin biopsy scores as reported ^54,55^.

### Cells culture and treatment

Human immortalized epidermal cells (HaCaT cells) were bought from iCell Bioscience Inc, Shanghai. Fibroblasts were obtained from skin tissues in accordance with a previous report ^56^. The cells were cultured in DMEM (GNM12800, Genom, Hangzhou, China) completed with 10% fetal bovine serum (FSP500, ExCell Bio, Shanghai, China) and 100 U/mL of penicillin, and 100 μg/mL of streptomycin (SV30010, HyClone, Logan, UT) in a humidified, 5% CO_2_incubator at 37 °C, and were harvested with a 0.25% trypsin/EDTA (25200072, Gibco, Grand Island, NY) solution.

For stimulation or inhibition assays, cells were cultured in medium containing MIF (100ng/ml, BK0133, bioworld, Nanjing, China) for 4h ^29^, ISO-1 (25μM, I303791, Aladdin, Shanghai, China) for 24h or 48h ^57,58^ and ZAKα-activating toxins anisomycin (ANS, 1μM, HY-18982, MCE) for 3h ^24^, pretreated with SB203580 (10 μM, S1863, Beyotime, Shanghai, China) for 2 h prior to UVB irradiation ^59,60^ and cultured in medium containing disulfiram (DSF, 40 μM) for 24h after UVB exposure ^61^ as previously reported.

### Cell transfection

HaCaT cells or primary mouse keratinocytes (iCell, Shanghai, China) were cultured in the 6-well plate to reach 60-80% confluence. Lipofectamine 2000 (11668-019, ThermoFisher) was used as a transfection reagent following the manufacturer’s instructions. After incubation for 8 hours at 37 ℃, the medium was removed and the cells were cultured in complete medium for another 24 hours for qPCR and 48 hours for WB. The sequences of siRNA (GenePharma, Suzhou, China) were listed in **Table S3**.

### Endogenous nucleic acids extraction and transfection

Cytosolic DNA and RNA were extracted from UVB-exposed HaCaT cells using DNA Extractor kit (BioTeke Corporation, Beijing, China) and EZ-press RNA Purification Kit (B0004DP, EZBioscience, Roseville, MN) following manufacturer’s instructions. Then the extracted eNAs (1.25 mg/ml DNA and 0.625 mg/ml RNA) were transfected to HaCaT cells using Lipofectamine 2000 as previous reported ^30^.

### AAVs production and injection

Lentiviruses were packaged in HEK293T cells using a three-plasmid packaging system consisting of packaging plasmid, helper plasmid, and Helper plasmid. The dsAAV serotype 2/9 (dsAAV2/9) was used in the present study. The regulatory element CMV was employed as a promoter, while the fluorescent labeled protein EGFP was utilized. The targeted sequence for Mif was CCGCAACTACAGTAAGCTG and the control sequence was CCTAAGGTTAAGTCGCCCTCG.

MRL/lpr mice were anesthetized and a subcutaneous injection was administered in the hair removal areas of the mice, consisting of a viral load of 1.0x10^11^ genomic copies of pscAAV-U6-shRNA (Mif)-CMV-EGFP-tWPA in a volume of 100µl. For the control group, an equal volume of pscAAV-U6-shRNA (NC2)-CMV-EGFP-tWPA was applied. One week after receiving the injection, the mice were utilized for the subsequent experiments.

### Fabrication and Characterization of Microneedle Patches

The microneedle patches were fabricated using a polydimethylsiloxane (PDMS) micromold, with the base length and the height of each microneedle measuring 350 μm and 850 μm, respectively. The needles were arranged in a 10 × 10 array with a needle pitch of 750 μm. Polyvinylpyrrolidone (PVP) solution was prepared and added with the ISO-1 (100 mM) or DMSO solution. The mixed solution was centrifuged for 20 min. Afterwards, the mixed liquid was injected into the microneedle mold and underwent overnight drying following vacuum treatment. Finally, the PVP solution was introduced into the microneedle mold and left to dry overnight, resulting in the formation of the patch base. The concentration of ISO-1 was approximately 235.4 μg per patch. After removed from the mold, the morphology of microneedles was determined using an optical microscope (Olympus CKX-41-32, Japan) and a scanning electron microscope (PHENOMPURE, Thermo Fisher, USA).

### Microneedle compression test and skin insertion test

The microneedles underwent the compression tests using a universal testing machine (PT-1699V, Baoda, China). The microneedle patch was placed in the center of the base with the needle tips facing up and then applied pressure vertically. The sensor-to-stage distance remained the same in all tests. The sensor was set to move at a rate of 0.2 mm/min until a displacement of 450 μm is reached. The trigger force was set to be 0.05 N, and the force-displacement curve of the microneedle was documented. For the skin insertion test, the patches were soaked with methylene blue solution and were pressed onto porcine skin with a pressure of 5-10 N for 5 min to assess the insertion capability. The inserted area was photographed using an optical microscope (Olympus CKX-41-32, Japan) after being torn off.

### Microneedle degradation test and skin irritation test

For the microneedle degradation test, the abdominal skin of the rats was shaved under anesthesia, and then the microneedle patch was firmly pressed on the abdominal skin and fixed with medical tapes. The patch was removed after 5 min, 10 min, 20 min and 30 min, respectively, and the inserted areas were photographed with an optical microscope (Olympus CKX-41-32, Japan). For the skin irritation test, the microneedle patch was pressed vertically into the abdominal skin and maintained for 30 minutes. Erythema and edema of the skins were documented at 0 hour, 1 hour, 6 hours, 12 hours, and 24 hours following removal.

### Immunohistochemistry

Sections of paraffin-embedded skin tissues were deparaffinized with xylene, rehydrated, and subjected to heat-induced epitope retrieval. Sections were blocked with goat serum, and incubated with a rabbit anti-human MIF, rabbit anti-human CD74 overnight at 4℃. Afterwards, sections were stained using a secondary antibody coupled with horseradish peroxidase (HRP), and the staining was visualized using the DAB kit (ZLI-9017, ZS, Beijing, China). The quantification was performed using Image-Pro Plus 6.0 (Media Cybernetics) as reported ^62^. The integrated optical density (IOD) is defined as the total count of positive materials across all locations within the region. The mean density in the selected area was calculated as IOD/Area. Information of antibodies is listed in Table S4.

### Immunofluorescence

Tissue sections were incubated in a mixture of two primary antibodies [rabbit anti-human/mouse MIF, mixed with mouse anti-human/mouse KRT14; rabbit anti-human CD74, mixed with mouse anti-human KRT14, or mouse anti-human vimentin] for overnight at 4℃. The sections were then incubated in a mixture of goat anti-mouse IgG H&L (Alexa Fluor ®488) and Goat Anti-Rabbit IgG H&L (Alexa Fluor® 555) in blocking buffer for 1 hour at room temperature. Sections were mounted using glycerol mounting media supplemented with anti-Fade with DAPI (ab188804, Abcam, Cambridge, UK). Slides were visualized using Fluorescence Microscopes (Olympus IX73, Japan). Information of antibodies is listed in **Table S4**.

### Quantitative real-time PCR

Total RNA was extracted from HaCaT cells, fibroblasts, as well as human and mouse skin tissues using EZ-press RNA Purification Kit (B0004DP, EZBioscience, Roseville, MN) according to the manufacturer’s instructions. Color Reverse Transcript Kit (A0010CGQ, EZBioscience, Roseville, MN) was used to synthesize cDNA. RT-qPCR was analyzed with Roche LightCycler® 480 instrument using SYBR Green Color qPCR Mix (A0012-R2, EZBioscience, Roseville, MN). The comparative threshold cycle (Ct) value of each sample was calculated, and the relative mRNA expression was normalized to the GAPDH or Actb value. Gene-specific primers are shown in **Table S5**.

### Western blot analysis

Whole-cell lysates were prepared with a radioimmunoprecipitation assay (RIPA) lysis buffer (CW2333S, CWBIO, Jiangsu, China) with protease and phosphatase inhibitors (P1045, Beyotime, Shanghai, China) for the total protein extraction. Protein concentration was determined by bicinchoninic acid (BCA) assay (23228, ThermoFisher). Primary antibodies (Anti-MIF; Anti-Collagen I; Anti-MMP2; Anti-MMP9; Anti-TNF alpha; Anti-IL-1 beta; Anti-MX1; Anti-p38; Anti-p-p38; anti-ZAKα; Anti-GSDMD; Anti-GAPDH) were used for immunoblot analysis according to the manufactures’ protocols. Phos-tag 7.5% SDS-PAGE gel (198-17981, Wako, Tokyo, Japan) was used for detecting phosphorylated ZAKα. Signals were observed using chemiluminescence ECL substrate (WBKLSO100, Merck millipore, Billerica, MA). The bands’ intensities were measured using ImageJ software. Details of the antibodies can be found in **Table S4**.

### ELISA assay

HaCaT cells were cultured in a 12-well plate until they reached 80%confluence. The medium was collected and the levels of MIF were quantified using Human MIF ELISA Kit (70-EK1158, MultiSciences, China) following the manufacturer’s instructions. The optical density (O.D.) was measured at 450 nm using an ELISA analytical instrument (Molecular Devices, SpectraMax® Plus 384).

### scRNA-seq analysis

The scRNA-seq data were acquired in GEO under accession number GSE186476. Data from a total of 14 normal skins and 7 pairs of lupus non-lesional (from sun-protected skin of the buttock) and lesional skins were included. The Seurat V3.0 R packages were utilized for analyzing, integrating, and normalizing datasets, as well as for performing dimensionality reduction, clustering, and differential expression of genes in the matrix. Cell clustering was performed using high variable genes, and the principal components based on high variable genes were used to generate the graphs at a resolution of 0.5.

### Statistical Analysis

All data were presented as mean ± SEM and analyzed using GraphPad Prism version 8.0 (GraphPad Software, San Diego, CA, USA). The student’s t-test was employed to compare two conditions, while one-way analysis of variance (ANOVA) was utilized to compare more than two conditions. A *P*-value less than 0.05 was considered statistically significant.

## Acknowledgements

Not applicable.

## Funding

This work was supported by the following grants:

National Natural Foundation of China Grant No. 81872524 (LW, Guangzhou), National Natural Foundation of China Grant No. 82073431 (LW, Guangzhou), National Natural Foundation of China Grant No. 81970632 (YZ, Guangzhou), Guangdong Science and Technology Department Grant No. 2020B1212060018 and 2020B1212030004 (YZ, Guangzhou).

## Author contributions

Chipeng Guo, Liangchun Wang, and Yiming Zhou designed the experiment. Liangchun Wang and Yiming Zhou funded the experiment. Chipeng Guo performed most of the *in vitro* and *in vivo* experiments, analyzed and generated data. Siweier Luo performed the bioinformatics analysis of scRNA-seq data. Tao Liu and Jigang Luo fabricated the microneedle patches. Haoran Lv, Yating Zhang, Le Wang, and Yi Xiao conducted the *in vitro* experiment and data collecting; Chipeng Guo and Yiming Zhou wrote the manuscript and all authors have revised and approved the manuscript.

## Competing interests

Authors declare that they have no competing interests.

## Data and materials availability

All of the data can be found in either the main text or the supplementary materials.

## Notes

### Competing Interest Statement

The authors have declared no competing interest.

