## Supplementary figures and images for "Ultraviolet B Induces Cutaneous Lupus Erythematosus by Triggering a Ribotoxic-Stress-Response-Dependent MIF-p38 Feedback Loop in Keratinocytes"

### Suppl. Fig.S1

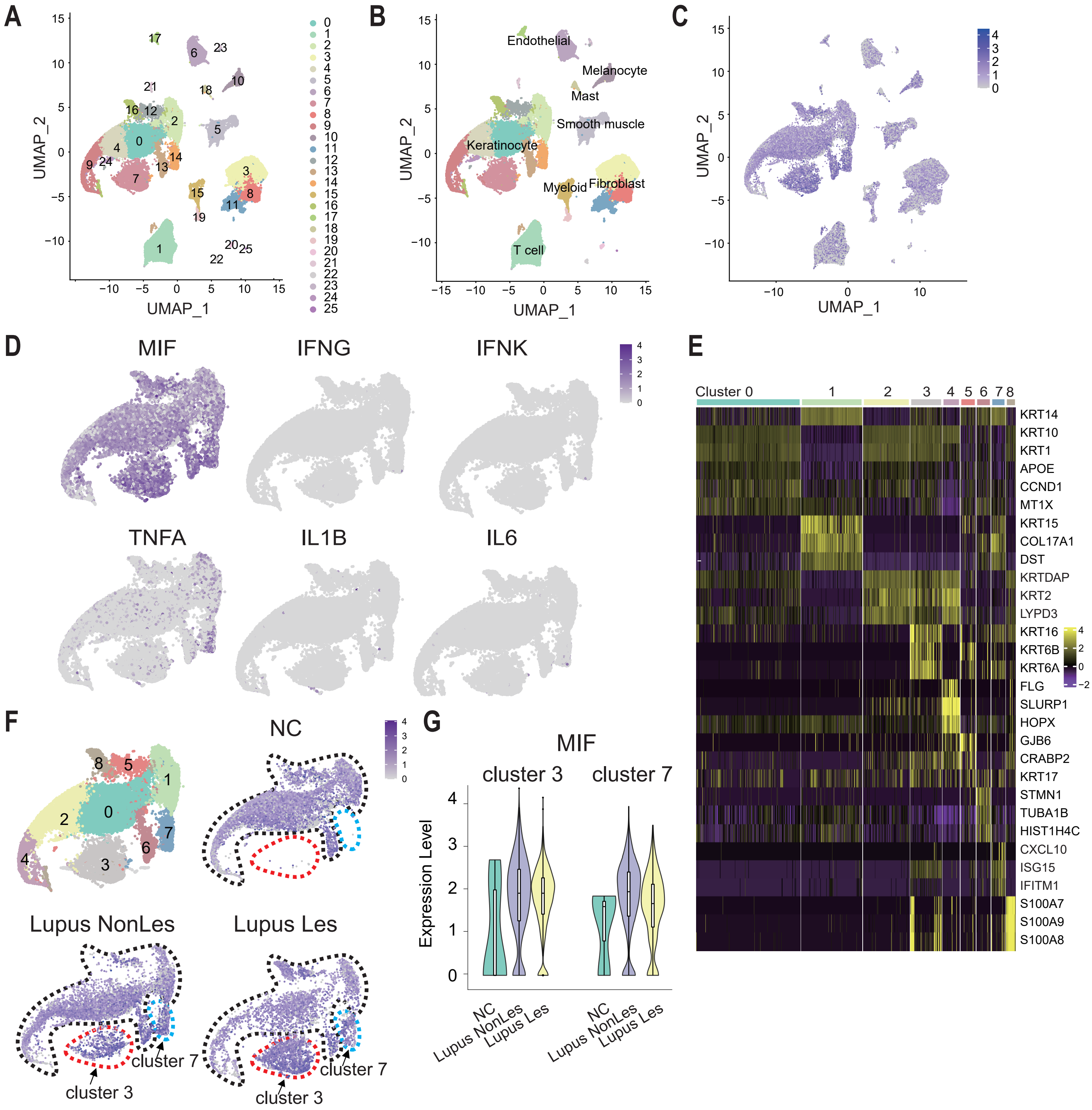

### Suppl. Fig.S2

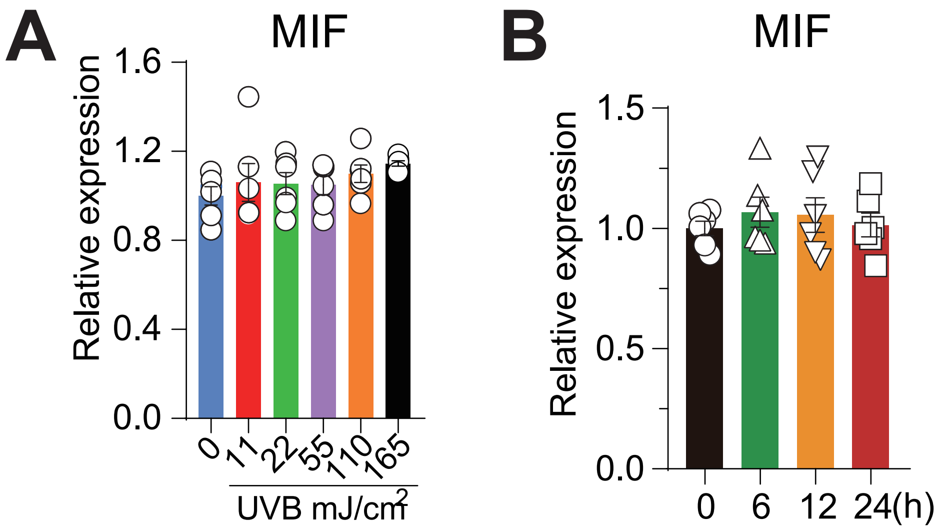

### Suppl. Fig.S3

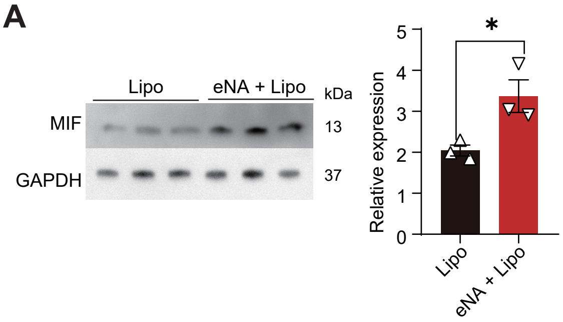

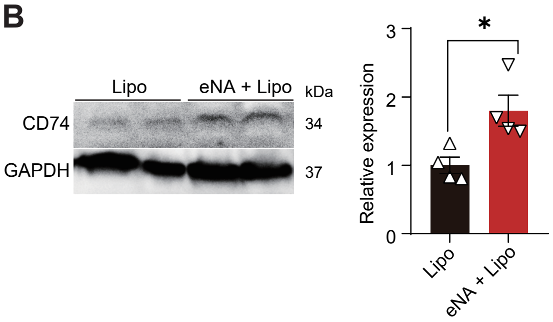

### Suppl. Fig.S4

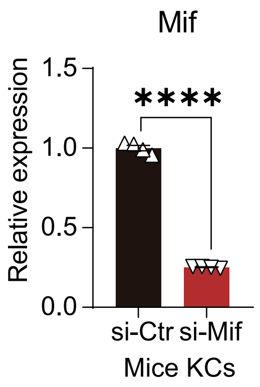

### Suppl. Fig.S5

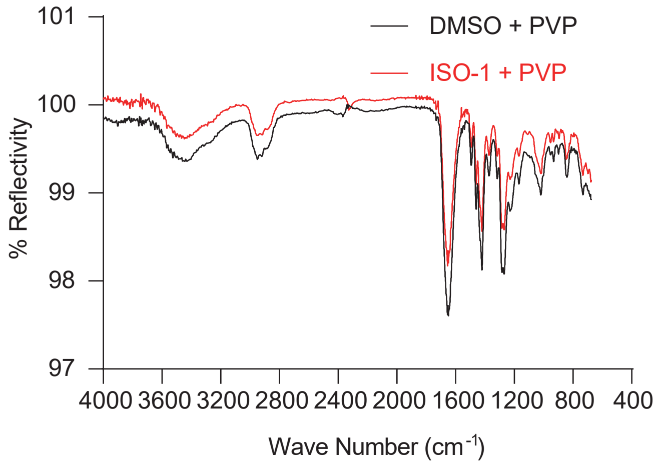

### Suppl. Fig.S6

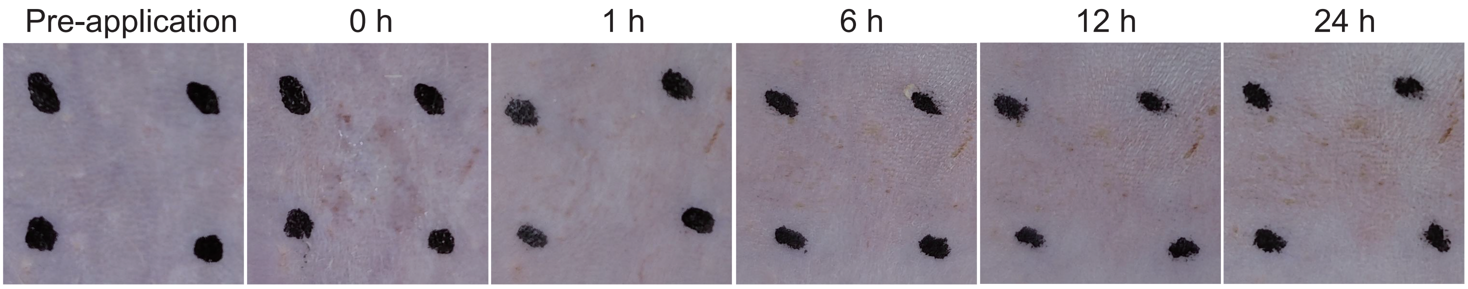
